# Effectiveness of an ultraviolet-C disinfection system for inactivating *Clostridioides difficile* spores in critical areas of a simulated patient room

**DOI:** 10.1101/2023.02.16.528885

**Authors:** Carolina Koutras, Richard L. Wade

## Abstract

*Clostridioides difficile* infections (CDI) pose a significant threat to patient safety in healthcare facilities. In this study, we investigated the effectiveness of an ultraviolet-C (UV-C) disinfection system in reducing *C. difficile* surface contamination in a simulated patient room.

The results showed an overall 98.17% reduction in *C. difficile* spores after UV-C treatment of directly exposed and shadowed areas, an efficiency that translated to a 4.91% reduction per minute of room vacancy. This study demonstrates that UV-C disinfection is an effective method for reducing *C. difficile* spores in healthcare settings, and that efficiency can be improved with shorter setup times and optimal device placement.

## Introduction

*Clostridioides difficile (C. difficile*) is a spore-forming, Gram-positive anaerobic bacillus. When *C. difficile* enters the body through the fecal-oral route, it can reach the large intestine (colon) and release exotoxins that destroy cells, cause inflammation, and trigger potentially lifethreatening diarrhea. *C. difficile* also forms highly resistant spores that easily persist in the environment and in the affected person, and continue to disseminate despite routine and terminal cleaning of hospital settings. C. difficile infections (CDI) are a significant concern for patient safety in healthcare settings. It has been suggested that the incidence of CDI in persons > 65 years of age contributed to over 80% of all deaths from CDI in recent years - an alarming and disproportionately high figure. [1,2,3] Thus, preventing CDI is a key aspect of patient safety in healthcare facilities. Here we investigate the effectiveness of an ultraviolet-C (UV-C) device in reducing *C. difficile* spores on high-touch surfaces in a simulated healthcare environment. Use of UV-C disinfection alone was proven effective to reduce 98.17% of all C. difficile spores on high-touch surfaces and shadowed areas of a mock patient room and bathroom. The rate of disinfection was 4.9% of spores per minute of room vacancy. UV-C is an effective infection control measure that can help hospitals reduce the risk of *C. difficile* spore exposure and protect patients from developing CDI.

## Methods

Using a modification of the ASTM International method E2197, spores of *C. difficile* (ATCC strain #43598) suspended in phosphate-buffered saline (PBS) with 5% bovine calf serum (BCS) were inoculated onto 20 mm stainless steel disks and dried. Disks were placed on right and left bedrails, under bed, call button, table top, table bottom, chair armrest, toilet seat, grab bar, sink handle, floor near device, floor far from device, in a simulated 3.0 × 5.8-meter hospital room with a 1.8 × 1.5-meter bathroom. A vertical 2.0 x 0.6-meter (height x width) ultraviolet-C tower equipped with 8 high output mercury lamps emitting at 254 nm (approximately 514 uW/cm2 total output at 2.5 meters) was used to treat the bathroom, and to treat each side of the patient bed (Arc, R-Zero). The bathroom was treated with a 6-minute cycle, with the tower placed 1.5 meters from and facing the sink faucet and 0.6 meters facing the center of the toilet. The patient room was treated with two 6-minute cycles, one on each side of the patient bed. In the first treatment position, the device was placed at 1.1 meters facing the center of the right side of the bed and 0.4 meters facing the center of the side table. The second positioning was also 1.1 meters facing the center of the patient bed left side and 0.4 meters from facing the center of the chair. On average, the device was placed 1.5 meters from all surrounding potential test locations, including bed rails, chair, table. After UV-C exposure, spores from exposed disks (three independent replicates per test location) and unexposed control disks were recovered in PBS with Triton X-100, enumerated using dilution plating, and log10 reductions were determined by comparing the number of spores recovered from exposed and control disks. The total number of tested carriers was 39, including controls. Times for set-up, treatment and resetting the space were recorded. Results are expressed as mean log10 reductions and percent reduction/minute of room vacancy (vacancy time due to UV-C treatment). A second test was conducted for comparison purposes and used the same parameters as above, except for UV-C exposure. During the second test, a 5-minute cycle was performed at the bathroom and on one side (right) of the patient bed.

## Results

The high-output UV-C tower tested achieved an overall 98.17% (1.74 log) reduction in *C. difficile* spores including both direct and indirect line of sight in the simulated patient suite (Table I). The locations closest to the device and within direct line of sight had the highest overall reductions (99.54-99.99% range). Even areas with no direct line of sight (shadowed) had an average of 93.83% reduction. All locations had a greater than 92.0 % reduction. The time required to set up the device for each of the three positionings was 10 seconds. The total time required to set up the device, including a built-in 30-second safety timer to allow time for the operator to safely exit the room, was 2 minutes for all three cycles combined. These overall efficiencies translate to percent reduction rates of 4.91% per minute of room vacancy (Table II). During a second test where only one cycle was performed in the bathroom and one in the patient room, for a total UV-C exposure time of 10 minutes as opposed to 18 minutes, an overall 87.99% reduction of *C. difficile* spores was observed.

**Figure 1.**
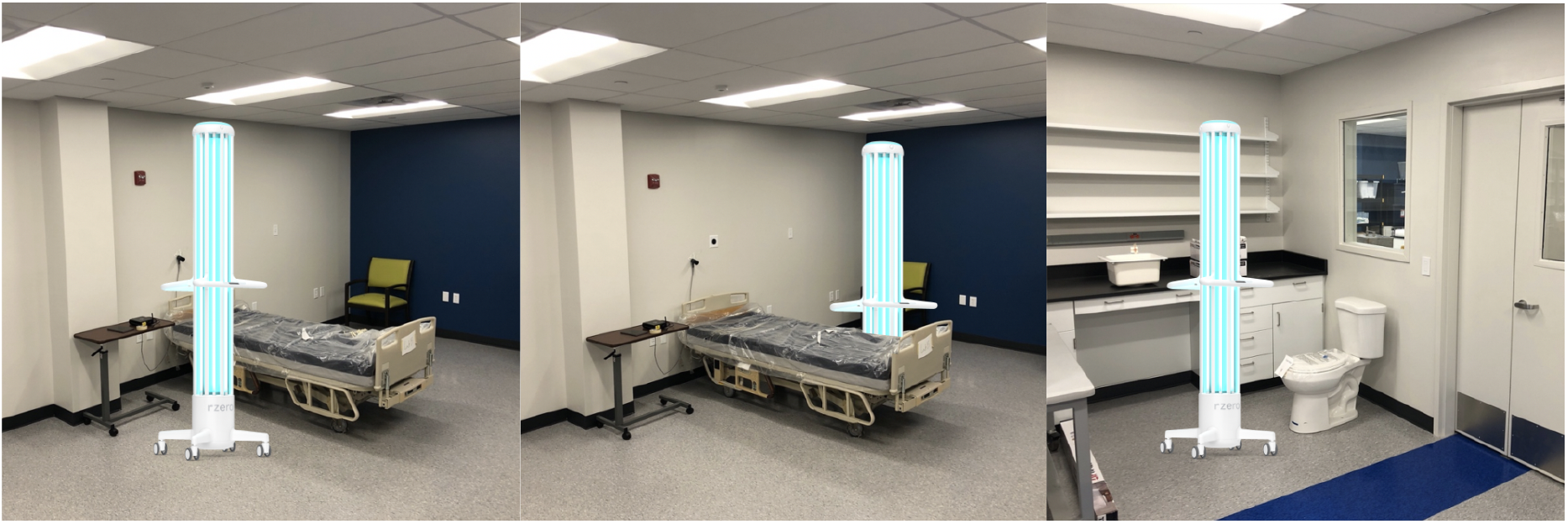
Study Diagram. The test device was positioned on both sides of the patient bed (first and second images to the left), and in the bathroom (right).

**Table I.** *C. difficile reduction* after UV-C treatment. The antimicrobial efficacy of the tested disinfection device is shown for all tested areas in a simulated patient room. The total UV-C run time was 18 minutes.

| Carrier | CFU/Carrier | Log Reduction (vs Control) | % Reduction (vs Control) |
| --- | --- | --- | --- |
| *Control* | 2.70E+06 | - | - |
| *Bedrail (right)* | 3.25E+02 | 2.51 | 99.99 |
| *Bedrail (left)* | 2.85E+04 | 4.45 | 98.94 |
| *Under Bed* | 2.00E+05 | 5.3 | 92.60 |
| *Call Button* | 1.37E+05 | 5.14 | 94.93 |
| *Guest Chair Armrest* | 4.03E+03 | 3.6 | 99.85 |
| *Floor (near)* | 2.78E+03 | 3.44 | 99.90 |
| *Table (top)* | 7.60E+03 | 3.88 | 99.72 |
| *Table (bottom)* | 1.34E+05 | 5.13 | 95.06 |
| *Floor (far)* | 3.92E+04 | 4.59 | 98.55 |
| *Toilet Seat* | 1.00E+03 | 3 | 99.96 |
| *Sink Handle* | 2.56E+04 | 4.41 | 99.05 |
| *Grab Bar* | 1.26E+04 | 4.1 | 99.54 |
| ***Overall % Reduction*** | **98.17** |  |  |

**Table II.**
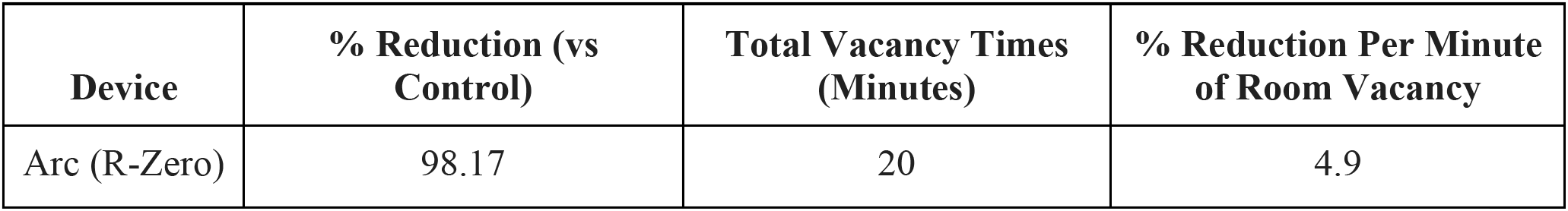
Device efficiency. The percent reduction per minute of room vacancy is shown below. The total vacancy time, including cycle setup times was 20 minutes.

| **Device** | **% Reduction (vs Control)** | **Total Vacancy Times (Minutes)** | **% Reduction Per Minute of Room Vacancy** |
| --- | --- | --- | --- |
| Arc (R-Zero) | 98.17 | 20 | 4.9 |

## Conclusion

There are many important factors that impact the efficiency of point-in-time UV-C disinfection devices in healthcare settings, including UV-C output, room configuration, operating procedures, vacancy times and percent reductions achieved during each minute of room vacancy. In this study, we tested the effectiveness of a high-output UV-C tower in reducing *C. difficile* in a simulated patient room. The tower was equipped with a safety timer and safety motion sensors, and had a manual setup time of only 10 seconds per cycle. A single 6-minute treatment cycle in both the bathroom and patient room areas resulted in nearly 88% reduction of all spores; overall disinfection efficacy was increased to 99.54% when an additional 6-minute cycle was incorporated into the protocol to maximize line of sight treatment of high touch surfaces on both sides of the patient bed. The data also supported the effectiveness of UV-C light on shadowed areas that are not easily touched. Chemical disinfectants constitute the first line of defense in infection prevention and control protocols. However, they do not eradicate *C. difficile* spores in healthcare, [4,5,6] where CDI is a significant concern for patient safety. Innovative UV-C towers are an important disinfection enhancement for patient rooms and beyond.

## Acknowledgements

The authors would like to thank Dr. Matthew Harwick and Resinnova Laboratories (Washington, DC) for their commitment to this project.

